# Speciation by Symbiosis: The Microbiome and Behavior

**DOI:** 10.1101/045195

**Authors:** J. Dylan Shropshire, Seth R. Bordenstein

## Abstract

Species are fundamental units of comparison in biology. The newly discovered importance and ubiquity of host-associated microorganisms is now stimulating work on the roles that microbes can play in animal speciation. We previously synthesized the literature and advanced concepts of speciation by symbiosis with notable attention to hybrid sterility and lethality. Here, we review recent studies and relevant data on microbes as players in host behavior and behavioral isolation, emphasizing the patterns seen in these analyses and highlighting areas worthy of additional exploration. We conclude that the role of microbial symbionts in behavior and speciation is gaining exciting traction, and the holobiont and hologenome concepts afford an evolving intellectual framework to promote research and intellectual exchange between disciplines such as behavior, microbiology, genetics, symbiosis and speciation. Given the increasing centrality of microbiology in macroscopic life, microbial symbiosis is arguably the most neglected aspect of animal and plant speciation, and studying it should yield a better understanding of the origin of species.

## MINIREVIEW

In 1998, Carl Woese referred to the microbial world as the “sleeping giant” of biology (1). Almost two decades later, unprecedented attention to our microbial world has turned the fields of zoology (2) and botany (3) inward - towards an increased awareness and understanding of individual animals and plants as holobionts (4–6). The term “holobiont” denotes a host plus all of its microbial symbionts, including inconstant and constant members that are either vertically or horizontally transmitted or environmentally acquired; it was first coined in 1991 by Lynn Margulis (reviewed in 5). The ubiquity and importance of microbes in and on holobionts, including humans, is evident in studies of host development (7), immunity (8), metabolism (9–12), behavior (13, 14), speciation (15, 16), and numerous other processes. Host-microbe interactions provide the holobiont with disadvantages (17–19) such as increasing the risk of cancer (20), and advantages (7, 21–23) such as driving the evolution of resistance to parasites and pathogens (24–26), and among other things producing signal components (i.e., metabolites) used to recognize differences in potential mates (27, 28).

The newfound importance of diverse microbial communities in and on animals and plants led to the development of the hologenome theory of evolution (4, 29). The “hologenome” refers to all of the genomes of the host and its microbial symbionts, and the theory emphasizes that holobionts are a level of phenotypic selection in which many phenotypes are produced by the host and microbial members of the holobiont. This developing scientific framework distinguishes itself by placing importance not only on well-studied primary microbial symbionts and vertical microbial transmission, but also on the vast diversity of host-associated microbes and horizontal microbial transmission. The key reason for aligning these different transmission modes and levels of complexity into an eco-evolutionary framework is that the community-level parameters among host and symbionts in the holobiont (e.g., community heritability, selection and coinheritance) can be analyzed under a common set of concepts to the parameters that occur in the nuclear genome (6, 30).

As natural selection operates on variation in phenotypes, the hologenome theory’s most significant utility is that it reclassifies the target of “individual” selection for many animals and plants traits to the holobiont community. This claim is straightforward given the overwhelming influence of microbes on host traits (31–34). The question going forward is whether the response to this community-level selection is relevant to the biology of holobionts. In other words, can host-associated microbial communities be selected such that shifts in the microbial consortia over multiple generations are a response to selection on holobiont traits? Community selection at the holobiont level is shaped by genetic variation in the host and microbial species and covariance between hosts and their microbial consortia, the latter of which can be driven by (i) inheritance of the microbial community from parents to offspring (35, 36) and/or (ii) community heritability *H^2^_C_* (30, 37). We recently summarized ten foundational principles of the holobiont and hologenome concepts, aligned them with pre-existing theories and frameworks in biology, and discussed critiques and questions to be answered by future research (6).

In the context of the widely accepted Biological Species Concept (38, 39), the principles of holobionts and hologenomes offer an integrated paradigm for the study of the origin of species. The Biological Species Concept operationally defines species as populations no longer capable of interbreeding. Reproductive isolation mechanisms that prevent interbreeding between holobiont populations are either prezygotic (occurring before fertilization) or postzygotic (occurring after fertilization). In the absence of reproductive isolation and population structure, unrestricted interbreeding between holobiont populations will homogenize populations of their genetic and microbial differences (6). While postzygotic isolation mechanisms include hybrid sterility or inviability, prezygotic isolation mechanisms can include biochemical mismatches between gametes and behavior mismatches between potential partners.

Symbionts can cause prezygotic reproductive isolation in two modes: broad-sense and narrow-sense (40). Broad-sense symbiont-induced reproductive isolation refers to divergence in host genes that result in a reproductive barrier because of selection on the host to accommodate microorganisms. In this case, loss or alteration of the symbiont does not have an impact on the capacity to interbreed; rather host genetic divergence and reproductive isolation evolve in response to microbial symbiosis and cause isolation regardless of whether the hosts are germ-free or not. Conversely, narrow-sense symbiont-induced reproductive isolation occurs when host-microbe or microbe-microbe associations result in a reproductive barrier, namely one that can be ameliorated or removed via elimination of the microbes. Therefore, narrow-sense isolation can be experimentally validated if it is reversible under microbe-free rearing conditions and inducible with the reintroduction of microbes. Isolation barriers that require host and microbial component underpin hologenomic speciation (6, 16).

We recently synthesized the literature and concepts of various speciation mechanisms related to symbiosis, with notable attention to postzygotic isolation (40–42). While aspects of the microbiology of prezygotic isolation are less understood, seminal cases exist (43–45) and control of behavior by symbionts is an emerging area of widespread interest (14, 46, 47). Here we emphasize the patterns seen in these new and old analyses (Table 1) and highlight important and tractable questions about the microbiome, behavior, and speciation by symbiosis. For the purposes of this review, we refer to the microbiome as the community of microorganisms in and on a host.

**Table 1.**
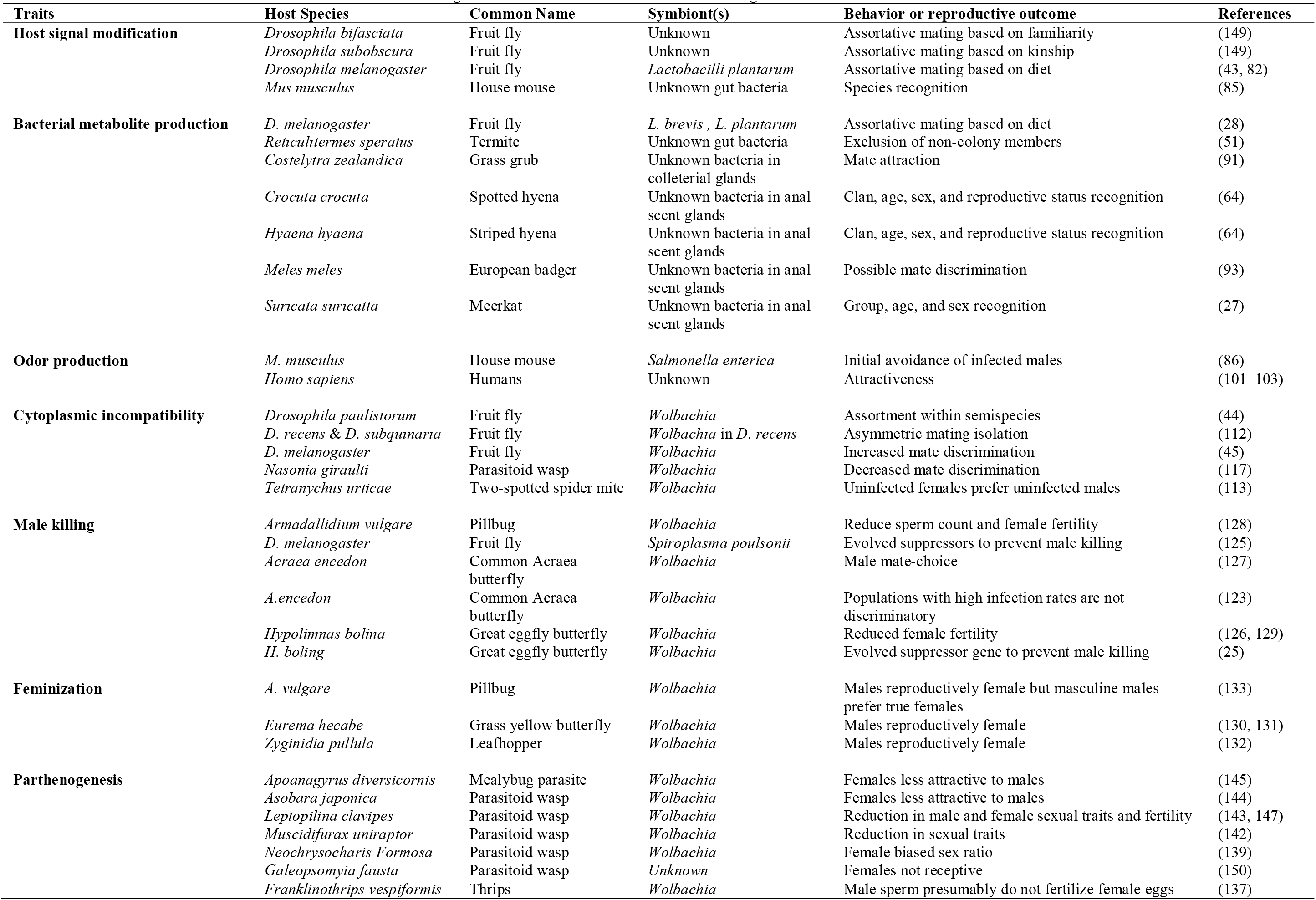
**Microbe-induced traits that associate with or cause changes in behavior and barriers to interbreeding**

## SIGNALING & MICROBIOME HOMOGENIZATION

Recognizing signals of species membership (48), gender (49), relatedness (50), and colony or group membership (51) is relevant to choosing a mate. Visual (48), auditory (49), and chemosensory signals (52) can each be used to relay this information, with the latter being particularly influenced by the microbiome in either “microbe-specific” or “microbe-assisted” ways. Both mechanisms involve the expression of chemosensory cues, but microbe-specific processes involve bacterial-derived products such as metabolites while microbe-assisted mechanisms involve bacterial modulation of host-derived odorous products (Fig. 1).

**Figure 1.**
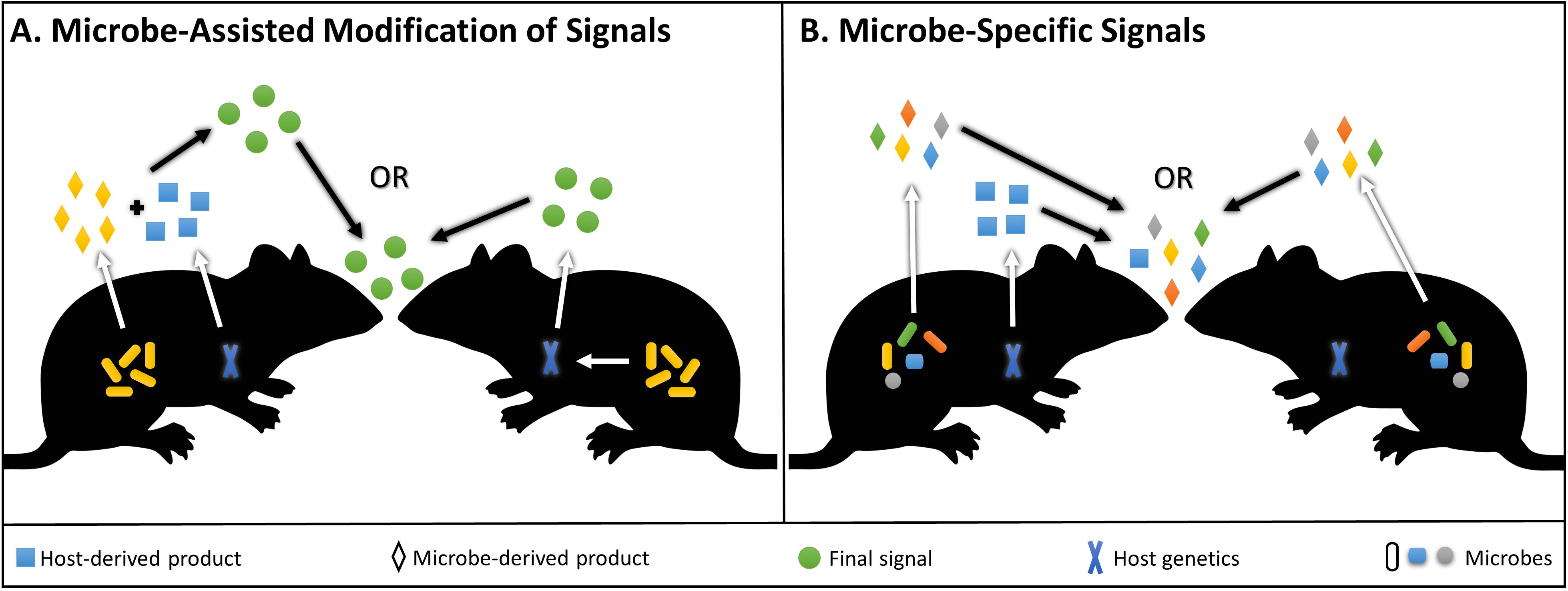
Microbe-assisted and microbe-specific signaling. (A) Microbe-assisted processes denote the production of a host signal with input from the microbiome. It occurs in two possible scenarios. On the left, the host and microbial symbionts produce products that interact or combine to form a signaling compound; on the right, microbial symbionts modify host signal expression, but they do not make a specific product directly involved in the signal itself. (B) Microbe-specific processes denote the production of a microbial signal without input from the host. It occurs in two possible scenarios. On the left, the host and microbial symbionts produce products that are both required to elicit a response; on the right, microbial symbionts produce compounds used by the host for signaling. Mouse image source: Wikimedia Commons, Angelus.

The microbiome’s capacity to provide identity information for mate recognition may rely on products being an honest signal of holobiont group membership, requiring that many or all members of the group (i.e., gender, population or species) contain appropriate microbial members that express equivalent signal profiles. Holobionts can be colonized by similar microbes via a number of different mechanisms, spanning behavioral similarities and contact with shared environmental sources (53, 54), similar ecological niches and diets (55–57), and host genetic effects (16, 58). Each of these mechanisms may explain a portion of the variation in the microbial communities of holobionts (40, 42, 59–61).

In the context of group living, humans in the same household (54, 62) and chimpanzees (63) or baboons (53) in the same social group have more similar microbial communities than non-group members. Among several mammalian species, microbial community composition covaries with odorous secretions, and similarities are shared based on host age, sex, and reproductive status allowing for potential signaling and recognition of these traits (27, 64). In hyenas, there is less microbial community variation within species than between them, and clans have more comparable microbial communities due to the marking and remarking of collective territory to signal clan ownership (64). In baboons, there is less microbiome variation within social groups than between them, and baboons involved in communal grooming behaviors share even more similarities (53). Insect populations such as termites can stabilize their gut microbiomes by way of trophallaxis, a behavior in which nestmates supply nutrients and microbes (e.g., cellulolytic microbes) to other colony members through fluids they excrete from their hindgut (65). However, Tung *et al* appropriately note, “*one of the most important unanswered questions is whether social network-mediated microbiome sharing produces net fitness benefits or costs for hosts*” (53). From the perspective of the origin of species, it will be similarly important to determine if fitness impacts of the microbiome in turn affect the evolution of group living and reproductive isolation. On one hand, socially-shared microbiomes could drive the evolution of population-specific mating signals and ensuing behavioral isolation. On the other hand, they could fuse incipient species in sympatry that socially share bacterial communities responsible for mating signals.

Similarities in diet can also influence microbiome homogenization, particularly in the digestive tract. For instance, *Drosophila melanogaster* reared on similar food sources carry comparable microbial communities (43). Trophically similar ant species also share microbial species (66). In humans, gut microbiome variation in taxonomy and functions correlates with dietary variation (67), and alterations in human diet can rapidly and reproducibly change the structure of the microbiome (68, 69). Seasonal variation in wild howler monkey diet is also correlated to shifts in the microbiome (70). Mediterranean fruit flies (71) and olive flies (72) acquire microbes from their food that increase clutch size and oviposition rate of females exposed to diets lacking essential amino acids (71, 72). Intriguingly, male sexual competitiveness of Mediterranean fruit flies increases up to two-fold with diets enriched with *Klebsiella ozytoca* versus a conventional diet (73).

Host genetics also affects microbial community assembly. In mice, there are 18 candidate loci for modulation and homeostatic maintenance of Bacteroidetes, Firmicutes, Rikenellaceae, and Provetellaceae in the gut (58, 74). Moreover, the presence of many rare bacterial groups in the gills of the Pacific oyster are correlated to genetic relatedness (75). Congruently, genetic variability in human immune-related pathways are associated with microbial profiles on several body sites including various locations along the digestive tract (76), and the largest twin cohort to date examined members of the gut microbiome and found that the bacterial family Christensenellaceae has the highest heritability (h^2^ = 0.39), and associates closely with other heritable gut bacterial families (77). Human genetic background also influences the risk of developing gastric cancer caused by *Helicobacter pylori*, indicating that incompatibilities between hosts and symbionts can produce deleterious effects (20). Phylosymbiosis, characterized by microbial community relationships that reflect host phylogeny (30), has also been reported in several cases. For instance, closely related *Nasonia* species that diverged roughly 400,000 years ago share more similar microbial communities than species pairs that diverged a million years ago (16, 40). Similar phylosymbiotic patterns are observed in hydra (59), ants (60) and primates (61).

The overall complexity inherent in microbial community structures and processes may be problematic for animal holobionts seeking to interpret a vast array of signaling information. However, recognition and differentiation of these microbe-induced signals may be possible if a subset of the microbiome affects the production of the particular signal. Furthermore, it may also be challenging to disentangle social, environmental, and diet effects on microbial assemblages in natural populations (53). Nonetheless, the important theme among all of these cases is that microbial community variation often appears to be less within holobiont groups/species than between them. This pattern, if sustained in natural populations, could facilitate the evolution of microbe-specific and/or microbe-assisted mating signals that promote recognition within populations or species and discrimination between them. Once this critical point is passed, speciation has commenced. There are parallels here with inclusive fitness theory, which posits that individuals can influence their own reproductive success or the reproductive success of other individuals with which they share genes (78, 79). If one follows the continuity from genes to microbial symbionts, then the inclusive fitness framework may also apply to holobionts in which specific microbial symbionts may influence their reproductive success by increasing the reproductive success of their hosts through microbe-specific and/or microbe-assisted mating. A case-by-case analysis of the reliance of the symbiont on the host for transmission (e.g., maternal, social, environmental transmission) will augment the relevance of this framework.

## MICROBE-ASSISTED MODIFICATION OF MATING SIGNALS

A common, microbe-assisted modification involves manipulation of host signals (Fig. 1A). One seminal study found that *D. melanogaster* acquires more *Lactobacillus* when reared on starch than on a molasses-cornmeal-yeast mixture (43, 80). The increased *Lactobacillus* colonization correlates with an upregulation of 7,11-heptacosadiene, a cuticular hydrocarbon sex pheromone in the female fly, resulting in an ability to distinguish fly holobionts raised in the starch environment from those reared on the molasses-cornmeal-yeast substrate (43, 81). This microbe-assisted positive assortative mating is reproducible, reversible, and maintained for several dozen generations after diet homogenization (43, 82). Moreover, this diet-dependent homogamy appears to be directly mediated by different gut bacteria, as inoculation of germ-free flies with *Lactobacillus* causes a significant increase in mating between flies reared on the different diets (43). Replication of these experiments found that inbred strains specifically followed this mating pattern (82). Moreover, another *D. melanogaster* study involving male mate choice and antibiotics revealed that female attractiveness is mediated by commensal microbes (83). These laboratory studies provide a critical model for how microbe-assisted modifications in a signaling pathway, ensuing behavioral changes, and mating assortment can potentiate behavioral isolation and possibly speciation. Indeed, natural populations of *D. melanogaster* express positive assortative mating and differential signal production based on food sources (84), and a bacterial role in these instances should be explored.

Microbe-assisted signaling also occurs in laboratory mice (*Mus musculus*), in which bacterial conversion of dietary choline into trimethylamine (TMA) leads to attraction of mice while also repelling rats (85). Antibiotic treatment decreases TMA production, and genetic knockout of the mouse receptor for TMA leads to decreased attraction in mice (85). Antibiotic treatment and subsequent depletion of TMA in mice could in turn result in a decrease in repellence of rats (85), though this possibility has not yet been tested *in vivo*. Another study found that female mice are more attracted to males not infected with *Salmonella enterica* infected compared to those that are, yet females mated multiply and equally in mating choice tests with the two types of males (86).

Mate preference based on infection status fits well with the Hamilton-Zuk hypothesis of parasite-mediated sexual selection, which posits that traits related to infection status can influence mating success (87). One seminal study showed that male jungle fowl infected with a parasitic roundworm produce less developed ornamentation and are less attractive to females (87). In house finches, male plumage brightness indicates their quality of broodcare and is associated with resistance to the bacterial pathogen *Mycoplasma gallicepticum* (88). The Hamilton-zuk hypothesis has been reviewed in detail (89).

## MICROBE-SPECIFIC SIGNALS

Microbe-specific signals frequently involve the release of volatile microbial metabolites, often through excretions from specialized glands on the host’s body (Fig. 1B). Microbial volatiles can transmit information utilized for social signaling (13, 90) and intra- or interspecies mate recognition (85, 91). For example, beetles (91), termites (51), nematodes (92), hyenas (64), meerkats (27), and badgers (93) produce and recognize bacterial metabolites in communication that can modulate their behavior. In termites, fecal metabolites produced by intestinal bacteria (51) coat the termite body and hive walls to signal colony membership. Termite holobionts lacking colony-specific metabolite profiles are attacked and killed by the hive (51). In contrast, some beetles and mammal species excrete bacterial metabolites from colleterial and anal scent glands, respectively (27, 64, 91). For example, female grass grub beetles house bacteria within their colleterial glands peripheral to the vagina that are used to attract males to mate (91).

An exciting area of research regarding microbe-specific bacterial signaling involves mammalian fermentation. The mammalian fermentation hypothesis (27, 64) states that fermentative bacteria within mammalian scent glands produce odorous metabolites involved in recognition. For example, hyena subcaudal scent pouches store bacteria that are mostly fermentative (64). When marking territory, hyenas deposit species-specific, bacterial-derived volatile fatty acids from this gland onto grass stalks (64). Bacterial metabolite secretions are more variable in the social hyena species, presumably because the complexity of signals from social species improves intraspecies identification (64). Alternatively, social hyenas may permissively transmit more diverse bacteria leading to diverse metabolite profiles. Hyena microbiomes also covary with group membership, sex, and reproductive state (64). Similarly, bacterial communities in meerkat anal scent secretions vary with host sex, age, and group membership (27). In both cases, the signal diversity may allow animal holobionts to recognize diverse biotic characteristics.

Humans also carry bacteria related to odor production. Breath (94, 95), foot (96), and underarm (97) odor covary with oral and skin microbiomes, respectively. Many diseases (e.g., smallpox, bacterial vaginosis, syphilis, etc.) are associated with distinct odors, and have historically been used by physicians in diagnosis (98). Clothing made from different materials even carry different odor profiles based on material-specific bacterial colonization (99, 100). Male odor has been associated with women’s interpretation of a male’s attractiveness (101–103), possibly influencing their choice in a mate.

The salient theme among the aforementioned cases is that host-associated microbes frequently emit odors, and sometimes this microbe-specific chemosensory information can affect mate choice. Reciprocally, ample evidence shows that chemical signals mediate sexual isolation (104), and a full understanding of whether these signals are traceable to host-associated microbes is worthy of serious attention. Germ-free experiments and microbial inoculations should be a prerequisite for such studies; otherwise they risk missing the significance of microbes in chemosensory speciation (104). Additional behaviors involved in speciation, such as habitat choice and pollinator attraction, are also likely to be influenced by microbe-specific products. Indeed, classic model systems of speciation await further experimentation in this light. For example, food-specific odors on apples and hawthorn translate directly into premating isolation of incipient host races of fruit flies of the genus *Rhagoletis* (105). Furthermore, the fruit fly *Drosophila sechellia* exclusively reproduces on the ripe fruit of *Morinda citrifolia*, which is toxic to other phylogenetically-related *Drosophila* species, including *D. melanogaster* and *D. simulans*. Some of the volatile compounds involved in these interactions, such as isoamyl acetate, have been associated with fermentative bacteria like *Lactobacillus plantarum* (106), suggesting that food-based premating isolation may be related to bacterial associations with the food source, though this requires further study. In summary, new challenges necessitate the concerted effort of scientists of diverse backgrounds to explore questions at the boundaries of many biological disciplines and to develop the tools to untangle and interpret this intricate web of interactions. Critical topics to be explored in the future include determining the microbial role in animal mate choice, quantifying the extent to which microbe-induced mating assortment impacts the origin of species, and identifying the mechanisms involved in these interactions.

## ENDOSYMBIONTS AND MATE CHOICE

*Wolbachia, Spiroplasma, Rickettsia, Cardinium*, and several other endosymbiotic bacteria can change animal sex ratios and sex determination mechanisms to increase their maternal transmission and thus frequency in the host population from one generation to the next. Notably, these reproductive alterations affect mate choice (107), and here we highlight a few prominent examples and discuss how endosymbiotic bacteria can influence behavioral isolation and the origin of species.

### Cytoplasmic Incompatibility

*Wolbachia* are the most well-studied reproductive distorters (108, 109) and are estimated to infect approximately 40% of all arthropod species (110). Across the major insect orders, *Wolbachia* cause cytoplasmic incompatibility (CI), a phenomenon in which *Wolbachia*-modified sperm from infected males leads to post-fertilization embryonic lethality in eggs from uninfected females or from females infected with a different strain of *Wolbachia*, but not in eggs from infected females (111).

In this context, *Wolbachia*-induced CI can promote the evolution of mate discrimination between populations or species because females can be selected to avoid males that they are not compatible with (Fig. 2C). Among closely related species of mushroom-feeding flies, *Wolbachia*-infected *Drosophila recens* and uninfected *D. subquinaria* contact each other and interspecifically mate in their sympatric range in Eastern Canada. However, gene flow between them in either cross direction is severely reduced due to the complementary action of CI and behavioral isolation. *Wolbachia*-induced CI appears to be the agent for evolution of behavioral isolation as asymmetric mate discrimination occurs in flies from the zones of sympatry but not in flies from the allopatric ranges (112). A similar pattern of *Wolbachia*-induced mate discrimination occurs among strains of the two-spotted spider mite, *Tetranychus urticae* (113) and *D. melanogaster* cage populations (45). Moreover, discrimination between particular semispecies of *D. paulistorum* is associated with their *Wolbachia* infections (44). In cases where host populations or species harbor different *Wolbachia* infections that are bidirectionally incompatible, for example in different *Nasonia* species that exist sympatrically (114, 115), reciprocal mate discrimination has evolved (114, 116). In contrast to these examples, interspecific mate discrimination in *Nasonia giraulti* is diminished when non-native transfections of *Wolbachia* spread throughout the whole body including to the brain, suggesting that *Wolbachia* can also inhibit pre-existing mate discrimination (117).

**Figure 2.**
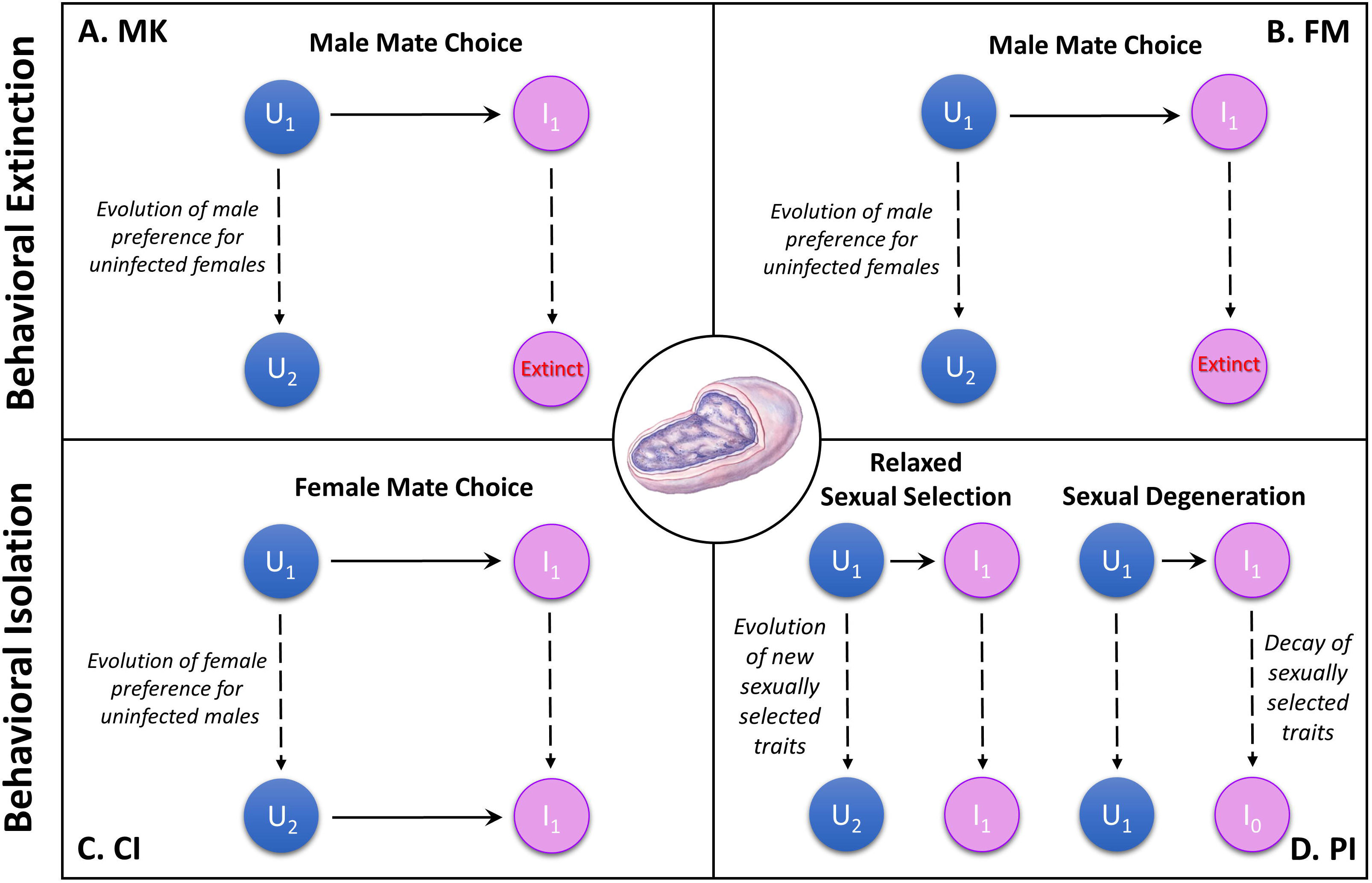
Endosymbiont-induced behavioral isolation and extinction. U (blue) and I (pink) represent the uninfected and infected populations, respectively. Horizontal solid arrows represent the direction of gene flow (from males to females) and vertical dashed arrows represent divergence time. Different subscript numbers for U and I represent evolutionary change in traits involved in behavioral extinction and behavioral isolation. Behavioral changes induced by (A) Male Killing (MK) and (B) Feminization (FM) evolve in response to selection on uninfected males to mate preferentially with uninfected females. If male preference is completely penetrant, then total loss of mating between the uninfected and infected population ensues, effectively leading the infected population to extinction since infected females rely on (the now discriminating) uninfected males to reproduce. We term this model “Behavioral Extinction”. In contrast, behavioral changes induced by (C) Cytoplasmic Incompatibility (CI) and (D) Parthenogenesis Induction (PI) can result in reduced or no gene flow between the infected and uninfected populations. CI-assisted reproductive isolation can be enhanced by the evolution of mate discrimination and specifically uninfected female mate choice for uninfected males. While this model does not sever gene flow in reciprocal cross directions, asymmetric isolation barriers can act as an initial step in speciation. PI-assisted reproductive isolation is mediated by two possible mechanisms: (i) Sexual Degeneration which involves the degeneration of sexual traits in the infected population that ultimately lock the populations into uninfected sexual and infected parthenogenetic species, and (ii) Relaxed Sexual Selection which involves the evolution of new sexual characteristics in the uninfected sexual population that prevent mating with the infected parthenogenetic population. *Wolbachia* image source: Tamara Clark, Encyclopedia of Life, *Wolbachia* page.

These cases reveal, to varying degrees, that *Wolbachia* can be causal to the evolution of assortative mating within and between species. Indeed, population genetic theory demonstrates that mate choice alleles spread quicker in populations or species with CI than those with nuclear incompatibilities (118). This is primarily due to the dominance of these *Wolbachia*-induced incompatibilities since CI causes F1 inviability, while nuclear incompatibilities are typically expressed in the F2 hybrids due to the recessive nature of hybrid incompatibility alleles.

### Male killing

Male killing is the most common form of endosymbiont-induced sex-ratio manipulation and can occur during embryonic (119, 120) or larval development (121, 122). The effect of male killing is to increase the number of female hosts in a population, thereby increasing endosymbiont transmission rates. To prevent complete fixation of females and population extinction (123), selection can favor hosts to (i) suppress male killing via genes that reduce *Wolbachia* densities or functions (25, 124–126) or (ii) electively choose mates whereby uninfected males preferentially mate with uninfected females (127, 128). If mate choice evolves as a behavioral adaptation to avoid male killing, it could begin to splinter infected and uninfected populations and initiate the first steps of the speciation process (Fig. 2A). One significant caveat in this conceptual model is that the infected population will go extinct without uninfected males to mate with. Thus, if mate preference based on infection status was complete, it would cause speciation between the infected and uninfected populations, resulting in the immediate extinction of the infected population that requires uninfected males to reproduce. We term this phenomena “behavioral extinction” (Figure 2).

*Wolbachia*-induced male killing can reach a state of equilibrium, as suggested by their long-term maintenance in natural populations of butterflies (129). Discriminatory males occasionally mate with infected females allowing for the infection to remain in the population (127), and eventually an equilibrium is reached (129). However in some cases, the infection rate is high (>95%), and male preference for uninfected females has not been identified (123). It is not known what mechanisms are involved in preventing male killing from reaching fixation in these situations.

### Feminization

Feminization, or the conversion of genetic males to morphological and functional females, has similar evolutionary consequences to male killing (Fig. 2B). This process occurs in many different arthropod species including butterflies (130, 131), leafhoppers (132), and woodlouse (133). Resistance to these effects in the pillbug *Armadillidium vulgare* has evolved in the form of feminization suppressors and male preference towards uninfected females. Males that mate with infected females produce feminized males (24, 134). Ultimately, a female-biased sex-ratio in feminized woodlouse populations results in an increase in male mate choice, male mating multiplicity, and sperm depletion. In the context of sperm depletion, initial mating encounters are normal, but upon increased mating frequency, males provide less sperm to subsequent females. Moreover, infected females are curiously less fertile at lower sperm densities possibly because they are less efficient at utilizing small quantities of sperm (128). Insufficient sperm utilization and slight differences in infected female courtship behaviors can result in male preference for uninfected females within the population (133). Just as with male killing, assortative mating within infected and uninfected populations may initiate the early stages of speciation and lead to behavioral extinction (Figure 2)

### Parthenogenesis

Microbial-induced parthenogenesis is common among haplodiploid arthropods such as wasps, mites, and thrips (135–137), wherein unfertilized eggs become females (138, 139). As we previously discussed (140), parthenogenesis-induced speciation by endosymbiotic bacteria falls neatly with the Biological Species Concept because parthenogenesis can sever gene flow and cause the evolution of reproductive isolation between sexual and asexual populations. Microbe-induced parthenogenesis does not necessarily exclude sexual capability of parthenogenetic females, but instead removes the necessity of sexual reproduction and can potentially drive divergence in sexual behaviors and mate choice (141). Speciation therefore commences between sexual and asexual populations under two models: (i) Sexual Degeneration and (ii) Relaxed Sexual Selection (140) (Fig. 2D).

The Sexual Degeneration model posits that the asexual population becomes incompetent to engage in sexual interactions due to mutational accumulation and thus trait degeneration while the sexual population remains otherwise the same (140). In this case, parthenogenetic lineages accumulate mutations in genes involved in sexual reproduction. Traits subject to mutational meltdown may span secondary sexual characteristics, fertilization, mating behavior, signal production, among others (142–144). For instance, long-term *Wolbachia-*induced parthenogenesis in mealybugs and some parasitoid wasps prevents females from attracting mates or properly expressing sexual behaviors (144, 145). Similarly in primarily asexual populations, male courtship behavior and sexual functionality is often impaired (142, 146, 147). The accrual of these mutations prevents sexual reproduction, thus causing the parthenogenetic population to become “locked in” to an asexual lifestyle. While this model is an attractive hypothesis for the onset of reproductive isolation between asexual and sexual populations, it is not always easily distinguishable from the alternative Relaxed Sexual Selection model (140). In this model, the sexual population diverges by evolving new or altered mating factors (e.g., courtship sequence, signals, etc.) while the asexual population does not degrade, but rather stays the same and thus can no longer mate with individuals from the diverging sexual population (140)

## CONCLUSIONS

Over the past decade, biology has stood *vis-à-vis* with what Carl Woese referred to as the “sleeping giant” of biology - the microbial world (1). During this period of groundbreaking research, a new vision for the increasing importance of microbiology in many subdisciplines of the life sciences has emerged. As such, studies of animal and plant speciation that do not account for the microbial world are incomplete. We currently know that microbes are involved in a multitude of host processes spanning behavior, metabolite production, reproduction, and immunity. Each of these processes can in theory or in practice cause mating assortment and commence population divergence, the evolution of reproductive isolation, and thus speciation. Understanding the contributions of microbes to behavior and speciation will require concerted efforts and exchanges among these biological disciplines, namely ones that embrace the recent “unified microbiome” proposal to merge disciplinary boundaries (148).

## ACKNOWLEDGMENTS

This publication was made possible by NSF grants DEB 1046149 and IOS 1456778 to SRB. JDS thanks Laci J. Baker for moral support during the writing process and Daniel LePage and Edward van Opstal for critique on figures. We also thank two anonymous reviewers for their critical feedback on the manuscript. Any opinions, conclusions or recommendations expressed in this material are those of the author(s) and do not necessarily reflect the views of the National Science Foundation.

